# Inferring Disease Risk Genes from Sequencing Data in Multiplex Pedigrees Through Sharing of Rare Variants

**DOI:** 10.1101/285874

**Authors:** Alexandre Bureau, Ferdouse Begum, Margaret A. Taub, Jacqueline Hetmanski, Margaret M. Parker, Hasan Albacha-Hejazi, Alan F. Scott, Jeffrey C. Murray, Mary L. Marazita, Joan E. Bailey-Wilson, Terri H. Beaty, Ingo Ruczinski

## Abstract

We previously demonstrated how sharing of rare variants (RVs) in distant affected relatives can be used to identify variants causing a complex and heterogeneous disease. This approach tested whether single RVs were shared by all sequenced affected family members. However, as with other study designs, joint analysis of several RVs (e.g. within genes) is sometimes required to obtain sufficient statistical power. Further, phenocopies can lead to false negatives for some causal RVs if complete sharing among affecteds is required. Here we extend our methodology (Rare Variant Sharing, RVS) to address these issues. Specifically, we introduce gene-based analyses, a partial sharing test based on RV sharing probabilities for subsets of affected relatives and an haplotype-based RV definition. RVS also has the desirable features of not requiring external estimates of variant frequency or control samples, provides functionality to assess and address violations of key assumptions, and is available as open source software for genome-wide analysis. Simulations including phenocopies, based on the families of an oral cleft study, revealed the partial and complete sharing versions of RVS achieved similar statistical power compared to alternative methods (RareIBD and the Gene-Based Segregation Test), and had superior power compared to the pedigree Variant Annotation, Analysis and Search Tool (pVAAST) linkage statistic. In studies of multiplex cleft families, analysis of rare single nucleotide variants in the exome of 151 affected relatives from 54 families revealed no significant excess sharing in any one gene, but highlighted different patterns of sharing revealed by the complete and partial sharing tests.

---

Sequencing distant relatives is an established approach to identify causal variants for Mendelian disorders [e.g., Ng et al. 2010, Bamshad et al. 2011, Ionita-Laza et al. 2011]. Typically external databases are combined with variant filtering strategies to identify causal variants under the assumption of complete penetrance and the absence of phenocopies. Sequencing related individuals has also become an option to identify causal variants in non-Mendelian complex disorders [Aida et al. 1998, Dahl et al. 2001, Daoud et al. 2009], although the rationale and strategies employed are more complicated. When familial phenotype aggregation is observed at a rate much higher than the prevalence in the general population, possible explanations include a shared familial environment or a relatively large familial “gene burden” [Isobe et al. 2013, Mescheriakova et al. 2016]. Another explanation could be the presence of a rare but highly penetrant (Mendelian or near-Mendelian) causal variant. This phenomenon has been observed in many common complex diseases and disorders such as chronic obstructive pulmonary disease, breast and ovarian cancer, birth defects, Alzheimer’s disease, and amyotrophic lateral sclerosis [Miki et al. 1994, Szabo and King 1995, Stratton 1996, Papassotiropoulos et al. 2006, Johnston et al. 2012, Bureau et al. 2014a].

We recently devised a statistical framework for such a setting, based on the notion that sequencing DNA in extended multiplex families can help to identify high penetrance causal variants too rare in the population to be detected through tests of association in population based studies, but co-segregating with disease within families [Bureau et al. 2014b]. Specifically, when only a few affected subjects per family are sequenced, evidence any one rare variant (RV) may be causal can be quantified from the probability of sharing alleles by all affected relatives, given it was seen in any one family member under the null hypothesis of complete absence of linkage and association. We presented a general framework for calculating such sharing probabilities when two or more affected subjects per family are sequenced, and showed how information from multiple families can be combined by calculating a p-value as the sum of the probabilities of sharing events at least as extreme [Bureau et al. 2014b]. We refer to this approach as RVS for Rare Variant Sharing. By sequencing three affected second cousins from a multiplex oral cleft family, we successfully employed this approach to identify a causal nonsense mutation in the gene *CDH1* [Bureau et al. 2014a].

Alternative approaches have also been proposed for the setting where the causal variant is rare, in the sense that when it is seen identical by state (IBS) in multiple affected relatives it has to be identical by descent (IBD). Instead of using exact sharing probabilities, Sul et al. [2016] proposed to use as test statistic the sum of the number of affected subjects sharing a RV and the number of unaffected subjects without the RV (either of which can be zero), standardized by subtracting its expectation and dividing by its standard deviation within each family under the null hypothesis and the assumption that only one founder in the family introduced a causal variant. Further, in both approaches inference is conditional on the introduction of the RV by a single founder, and as a consequence we do not need to know or estimate its population allele frequency. This is a great benefit, as the allele frequency is commonly unknown, especially when sequenced families come from a genetic background not well represented in the standard reference databases such as the Exome Sequencing Project [Fu et al. 2013] or the 1000 Genomes Projects [1000 Genomes Project Consortium et al. 2015], or the sample is a mix of families from different genetic backgrounds. Qiao et al. [2017] proposed GESE, a gene-based segregation test requiring an estimate of variant frequencies (as compared to calculating the probability of sharing conditional on the variant being observed), but otherwise relying on very similar assumptions as our previously introduced RVS approach [Bureau et al. 2014a]. Specifically, GESE (like RVS and RareIBD) assumes only one founder in the family introduced a causal variant in a gene, and Qiao et al. [2017] recommend limiting the tests to variants with high functional impact. Further, GESE also calculates the p-value as the sum of the probabilities of all events as or less likely as the observed event. However, in addition to absence of phenocopies (i.e., all affected subjects in a family are carriers), GESE assumes complete penetrance (i.e., all unaffected subjects do not carry the putative causal variant), while our original variant sharing approach was based only on sharing among affected subjects, and did not make any assumptions about unaffected subjects. The latter feature can be critically important, as was the case in detection of a nonsense mutation in *CDH1* shared among three affected second cousins, but also present in the unaffected parents who transmitted this variant [Bureau et al. 2014a, Figure 1]. We note GESE and RareIBD can be applied in “affected-only” mode by setting the phenotype of unaffected subjects to unknown, even though not intended as such.

**Figure 1:**
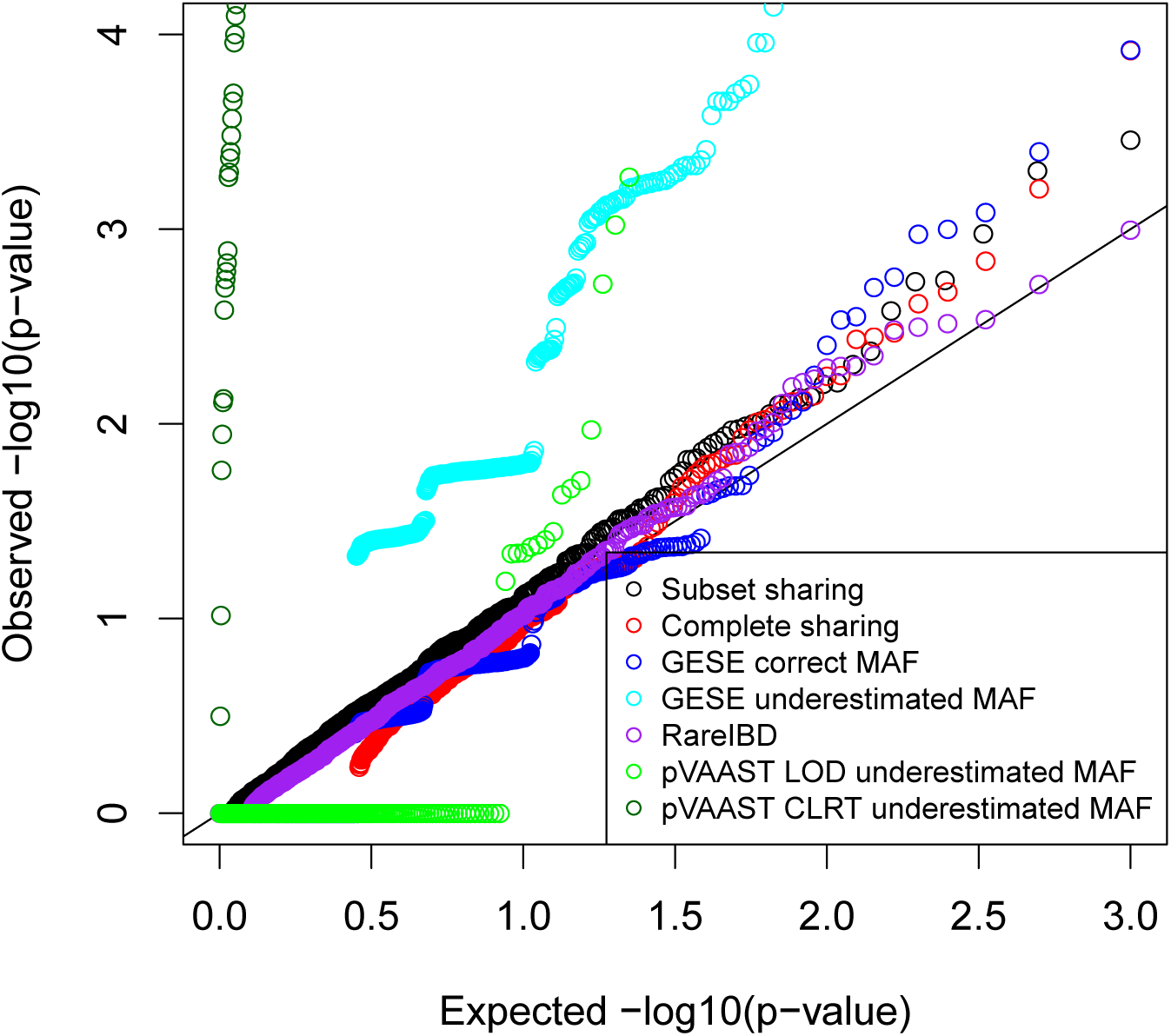
Expected and observed p-values under the null hypothesis in 1000 replicates (200 for pVAAST).

When reliable information about allele frequencies is available, some authors have argued for combining a linkage signal with the association signal derived using known allele frequencies to increase statistical power. Hu et al. [2014] proposed an extension of their previously developed Variant Annotation, Analysis and Search Tool (VAAST), a likelihood-ratio-based RV association test that incorporates case/control allele frequency differences and functional annotation into the likelihood. The aptly named pedigree-VAAST (pVAAST) employs a sequence-based model, where variants are tested for being directly causal instead of merely linked to some unobserved disease variant (as in classical linkage analysis) in variant and gene-based linkage analysis, and offers the option to combine this linkage information with the VAAST statistic. GESE calculates a probability of segregation combining association and linkage signals requiring knowledge (or an accurate estimate) of the variant frequency in the population. This feature of GESE has the potential to increase power, but misspecification of variant frequencies, which is likely in family samples with distinctly different or mixed genetic backgrounds, may yield spurious signals. Methods relying on filtering of variants based on frequencies in external databases such as our RVS approach and RareIBD are also susceptible to false positive signals from variants common in a study population but not well represented in the utilized reference databases. However, the haplotype structure around a variant contains information on the population frequency of variant that can be exploited to filter out common variants without producing a frequency estimate as required by pVAAST and GESE.

GESE, pVAAST and RareIBD are designed for analysis of all RVs in a gene or genomic region, as is standard when analyzing RVs to increase the proportion of subjects or families with at least one RV [Li and Leal 2008, Lee et al. 2014]. Here we present an extension of our previously published single-variant RVS method for a gene-based approach. Acknowledging phenocopies, diagnosis error and intra-familial genetic heterogeneity exist in complex disorders, we further extend our method by relaxing the previous assumption that all sequenced affected subjects must be carriers of the same causal variant, and by introducing an approach based on haplotypes of known variants to detect which variants are not actually rare in the study population despite being rare or completely absent in reference databases. Comparing our gene-based RV sharing approach to alternative methods, we show in a simulation study that knowledge and use of allele frequencies of rare variants in approaches such as GESE and the pVAAST linkage statistic does not lead to power gains over methods such as RVS and RareIBD, which do not require knowledge of such frequencies and are therefore more universally applicable. An implementation of our method RVS is available as open source software from the Bioconductor project at bioconductor.org/packages/release/bioc/html/RVS.html.

## Methods

### Gene-based analysis

We initially presented the RVS approach for single variants [Bureau et al. 2014b]. As long as there is a single RV within a gene in the same family, the RVs are independent since the families are unrelated, and the information can simply be pooled together and analyzed jointly. The abundance of RVs in the human genome, however, implies that multiple RVs are likely to occur on the same haplotype over a region such as a gene. Such RVs have identical sharing patterns, and are indistinguishable in genetic analysis. Therefore, we redefine the units of analysis as the haplotypes of RVs over each genomic region instead of individual RVs themselves. Simply taking the minimal RV sharing probability among all RVs in the same gene in a family has the effect of merging RVs on the haplotype with the lowest sharing probability. We detail in the section “Recoding rare variants haplotypes for rare variant sharing computations” on page 7 a systematic approach to recode RVs into haplotypes based on genomic sequence. After this recoding, when two or more RVs (or haplotypes of several RVs) remain in the same gene in a family, we retain one RV per family to compute the RV sharing probability. We propose using those RVs with the sharing pattern yielding the lowest probability among all RVs present in the same gene. When we do this, the test is no longer exact; the resulting p-value becomes an approximation of the exact p-value. We examine the impact of this practice on Type I error in a simulation study, and in sequencing data from individuals drawn from multiplex cleft families.

### Partial sharing

We define the following random variables:

*C_i_* represents the number of copies of the RV observed in the sequence of subject *i*,

*F_j_* is the indicator variable that founder *j* introduced one copy of the RV into the pedigree, and

*D_ij_* is the number of generations (meioses) between subject *i* and his or her ancestor *j*.

For a set of *n* sequenced subjects for which the pedigree structure limits to one the number of copies of the RV that they can share, we compute the probability that any subset of size *k* ≤ *n* shares a given RV. Without loss of generality, we assume the *n* subjects are ordered such that subjects 1, …, *k* ≤ *n* share the RV, and thus:

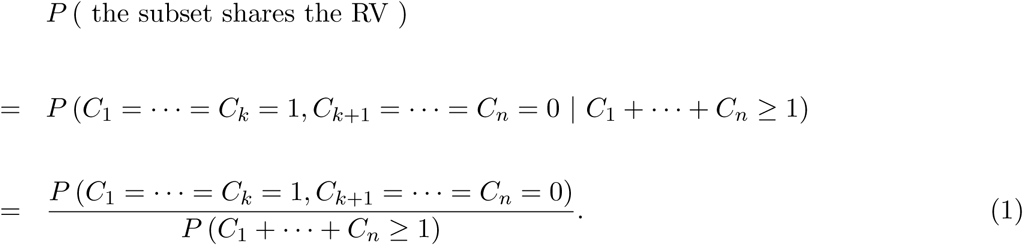

To simplify the notation, we define *K* = *C*_1_ + ⋯ + *C_n_*. For a single family, we define a RV sharing configuration where *k* subjects share a RV as *G_k_* = (*C*_1_,…*C_n_*) | *K* = *k*. The p-value of this configuration *G_k_* is the sum of probabilities of all sharing configurations *g* with probability *P_g_* = *P* (*g* | *K* ≥ 1) lower or equal to the probability of the observed configuration 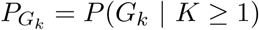, and with size *k*^(^*^g^*^)^ ≥ *k*, i.e.:

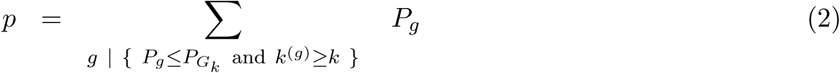

We note it is computationally advantageous to identify classes of all equiprobable configurations, to compute their probability only once for a member of each class, and to multiply the probability by the number of equiprobable configurations. Classes of equiprobable configurations are defined by exchangeable relatives (e.g. siblings), and subsets of exchangeable relatives (e.g. sibships who represent sets of first cousins).

With *M* families where the same RV is observed, or where *M* distinct RVs are observed (a different one in each family) in the same region (such as a gene), the configuration 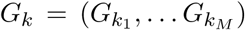 is a vector of family-specific configurations containing the *k*_1_, …, *k_M_* subjects sharing a RV in the *M* families, with 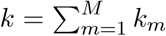. The p-value is then computed by applying the same criteria used for a single family, with the only modification to the probability of a global configuration *g* = (*g*_1_,…*g_M_*) with 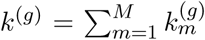 affected subjects being the product over the probabilities of all family-specific configurations *g*_1_,…*g_M_* under the independence assumption:

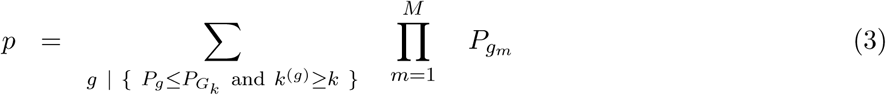

In the current implementation, *M* is limited to 10 as the summation grows exponentially with *M*. This *M* is the number of families with a RV, not the maximum size of the family sample, as only a fraction of families harbor RVs in any given gene.

### Refining the definition of RVs

The assumption that all copies of a RV seen in related members of a pedigree are IBD is crucial to the validity of the RVS tests. Instead of relying solely on filtering of common variants based on their frequencies in external reference databases, genome-wide genotype data enables the identification of variants actually introduced multiple times to the family from unrelated founders and most likely not rare in the population from which the family is drawn. Our approach is based on haplotypes of known variants (common and rare). RVs seen on two or more haplotypes are discarded as introduced multiple times into the family, e.g. being IBS without being IBD. We infer haplotypes by phasing the sequence data in families prior to analysis. Given the need to include a sufficient number of common variants within a genomic interval spanning a gene, and to ensure the quality of phasing, this approach is advisable only with genomic sequence and not with capture-based exome sequencing data alone. Our implementation uses Shapeit2 available at mathgen.stats.ox.ac.uk/genetics_software/shapeit/shapeit.html, with the duoHMM option to improve phasing where parent-offspring pairs are sequenced [O’Connell et al. 2014]. Haplotypes for the purpose of determining IBD status are comprised of variants previously observed in genomic sequence databases (such as the 1000 Genomes project), as these variants are less likely to be sequencing errors. They may include RVs included in the analysis, depending on the filter used to define a RV. We have opted to include all variants within the RefSeq gene boundaries in the inferred haplotypes.

### Recoding rare variant haplotypes for rare variant sharing computations

Once RVs have been assigned to haplotypes, those haplotypes containing at least one RV meeting the filtering criteria are recoded as a new synthetic RV representing all the RVs on that haplotype. Hence, there are as many recoded RVs in the analysis region as there are different haplotypes bearing at least one RV. The RV sharing probabilities are then computed for each recoded RV. In the same spirit, Sul et al. [2016] considered variants with the same genotype in all family members as duplicates and used one of those variants in the computation of the RareIBD statistic.

### Implementation

Our RVS approach is freely available through the `RVS` Bioconductor package [Sherman et al. 2018]. Briefly, the main function `RVgene` takes as input a data frame with pedigree and genotype data in standard formats (two alleles of a variant on two consecutive columns or minor allele count on one column) In addition, lists of RV sharing probabilities (pre-computed by the `RVsharing` function) and numbers of affected subjects for each possible sharing configuration of the sequenced affected members from every family must be provided to the `RVgene` function to perform the test allowing for partial sharing. Due to the computational demand of the convolution of the RV sharing event distribution for all families involved, the number of families with RVs in the same gene is currently limited to 10, or fewer depending on available RAM (the complete sharing test does not have this limitation). Another new feature of the `RVS` package is its correction of sharing probabilities to account for cryptic relatedness using the analytical approximation described in Bureau et al. [2014b], based on a reimplementation of the RV sharing probability computation using the `gRain` package for general computations on Bayesian networks. The previous Monte Carlo approximation for cryptic relatedness correction has also been reimplemented.

### Simulation study

We used 594 phased sequences from the CEU, TSI and GBR samples of the 1000 Genomes Project to define a population of haplotypes for each gene in the genome. Single nucleotide variants (SNVs) seen more than once in this sample were used to define recoding haplotypes. RVs were defined as SNVs with frequency < 1% among the 594 sequences (up to 5 copies). To evaluate haplotype recoding in the context of a sequencing study with a family sample, we assumed a genetic origin to the disease in all families and used the 47 simple pedigree structures (i.e., not allowing inbreeding or marriage loops, removing 7 inbred pedigrees to limit computational complexity) from the second set of multiplex cleft families (see section “Multiplex oral cleft families” on page 10). To evaluate statistical properties of this test, focusing on distant affected relatives for which the RVS tests are designed, we further removed the 14 pedigrees with affected 1^st^ degree relatives, leaving 33 simple pedigrees including 93 affected 2^nd^ – 9^th^ degree relatives. We assumed the disease had a genetic origin in 16 families (randomly sampled at each replicate) or about 50% of all available families. For each RefSeq gene, each family where the disease had a genetic origin was assigned a RV as potential disease causing variant (a distinct RV for each family if there were enough RVs in the gene, otherwise RVs were reused across multiple families). The genotype at the causal RV site was generated conditional on disease status under the Risch 2-locus heterogeneity model of disease [Risch 1990, and Table 1], which means the potential causal RV was not necessarily present in the family (the disease could be caused by the other unlinked gene). The disease prevalence implied by this model was 2.5%. Founder haplotypes were then sampled from the population of all haplotypes, and transmission to descendants was simulated without recombination. Both haplotype sampling and transmission were conditional on genotypes at the causal RV site. In families where the disease did not have a genetic origin (a subset of families for power evaluations, all families for assessment of Type I error under the null hypothesis), founder haplotype sampling was performed unconditionally, and transmission was simulated under Mendelian segregation instead.

We compared the statistical power of gene-based tests including the proposed RVS test allowing for partial sharing, the original RVS test (complete sharing), RareIBD [Sul et al. 2016], as well as two competing tests requiring external variant frequency estimates, namely GESE [Qiao et al. 2017] and the pVAAST LOD score [Hu et al. 2014]. We define RVs as variants with a minor allele frequency (MAF) less than 1%, and evaluated power for a gene with few coding RVs (*PEAR1*, 7 coding RVs) and a gene with many coding RVs (*CDH13*, 20 coding RVs). Separate simulation using RV penetrances of 0.25, 0.5 and 0.75 were conducted, corresponding to relative risks of about 10, 20 or 30 under the respective genetic models (Table 1). We also assessed Type I error of the gene-based tests in the context of a gene with many rare coding RVs (*CDH13*), where there were usually multiple RV-carrying haplotypes in the same gene in the same family.

**Table 1:** Model parameters used in the simulations (MAF: minor allele frequency).

| Locus | MAF | Penetrance |
| --- | --- | --- |
| Tested | $10^{-4}$ | $x \in \{0.25, 0.50, 0.75\}$ |
| Other | 0.1 | 0.04 |
| None (phenocopy) | NA | 0.02 |

Both GESE (in the case where no external estimates are available, for example from public databases) and pVAAST require control samples, so we created samples of 5,000 control subjects by randomly sampling two haplotypes for each subject from the 594 phased 1000 Genome sequences. This is the minimal size recommended by Qiao et al. [2017] to ensure a correct Type I error for GESE. Variants with MAF less than 1% in the control data were selected. For pVAAST, we specified a genetic model where the tested RV was assumed to be dominant with a penetrance in the range 0.5 – 1.0 (the interval over which pVAAST maximizes the LOD score). Since pVAAST requires specifying the amino acid substitution at the tested RV, we assumed a substitution with a mild effect (alanine to valine). The severity of the mutation is actually irrelevant for the linkage LOD score statistic, but is taken into account in the case-control composite likelihood ratio test (CLRT) which is added to the linkage LOD score to obtain the CLRT for pedigrees (CLRTp). Due to high computational demands of pVAAST, the number of Monte Carlo simulations to estimate the p-value was set to 10,000. For RareIBD and GESE, the number of resampling simulations was adaptively selected, increasing up to 10^7^ for very small p-values. For GESE, we evaluated the Type I error with control samples drawn from a population with the correct MAF and a population where the true MAF of the RVs was 10 times lower to assess the impact of MAF misspecification. Again, due to excessive computational burden, assessment of the Type I error for pVAAST was limited to 200 replicates and was performed only under the latter control sample definition.

### Multiplex oral cleft families

The first set of 55 multiplex cleft families was described in detail in Bureau et al. [2014a]. In brief, the families were mostly comprised of pairs of distant affected relatives with a few triples of distant affected relatives, on which whole exome sequencing (WES) was performed. We refer to this sample as the “WES sample”. Two of the multiplex families in the WES sample were expanded to additional affected relatives, and included in the second set of families described below. For that reason, these two families were removed from the WES sample in the current analysis, leaving 53 multiplex families. The second set included 54 multiplex cleft families from the Philippines, the United States (European ancestry), Guatemala and the Syrian Arab Republic. Whole genome sequencing (WGS) data were generated for 153 affected relatives and 7 unaffected relatives (all unaffected individuals were from Filipino families). We thus call this set the “WGS sample”. In one Syrian pedigree, the 8 affected relatives did not have any known common ancestor, so a sub-pedigree including 6 affected relatives descending from a common couple of ancestors was used in the analysis, reducing the total number of affected subjects to 151 (Table 2). The sequencing, alignment, and variant calling process was described in Holzinger et al. [2017], who also reported on RVs observed in the WGS data from the Filipino and Syrian families.

In addition to a large proportion of families with four or more affected relatives, the WGS sample differs from the WES sample by the presence of first degree relatives (Supplementary Table 1). Thus, the distribution of the − log_10_ probabilities of sharing a RV by all affected relatives within the respective families (which is the potential − log_10_ p-values when a RV is present in a single family) was more dispersed in the WGS than the WES sample (Supplementary Figure 4).

SNVs from both WES and WGS data were annotated using wAnnovar (wannovar.wglab.org) in November 2016. Exonic and splice site SNVs were extracted and filtered using the same criteria as described in Bureau et al. [2014b], with the additional requirement of a maximum frequency of 1% in all population samples from the gnomAD exome database (gnomad.broadinstitute.org), but dropping the previous step of filtering against the internal database of the Center for Inherited Disease Research. For the WGS data the above described haplotype-based approach was applied to ensure rare SNVs were introduced only once in each family. The duoHMM algorithm of Shapeit2 [O’Connell et al. 2014] made use of 37 parent-child duos of sequenced subjects to improve phasing, while the remaining 112 sequenced subjects were treated as unrelated (more distant sequenced relatives are not considered by the duoHMM algorithm). In the WES data, haplotyping of SNVs on a SNP array (Illumina Omni Express) with Merlin was used to infer IBD sharing from common variants, but did not use rare SNVs. Gene-level p-values were computed for both partial and complete sharing tests, separately for the WES and WGS samples, using the above described approach. Genome-wide association study results in the top genes detected by the RVS analysis were extracted from the Facebase Human Genomics Analysis Interface, Pittsburgh, PA (http://facebase.sdmgenetics.pitt.edu) [accessed June 20, 2018].

**Table 2:**
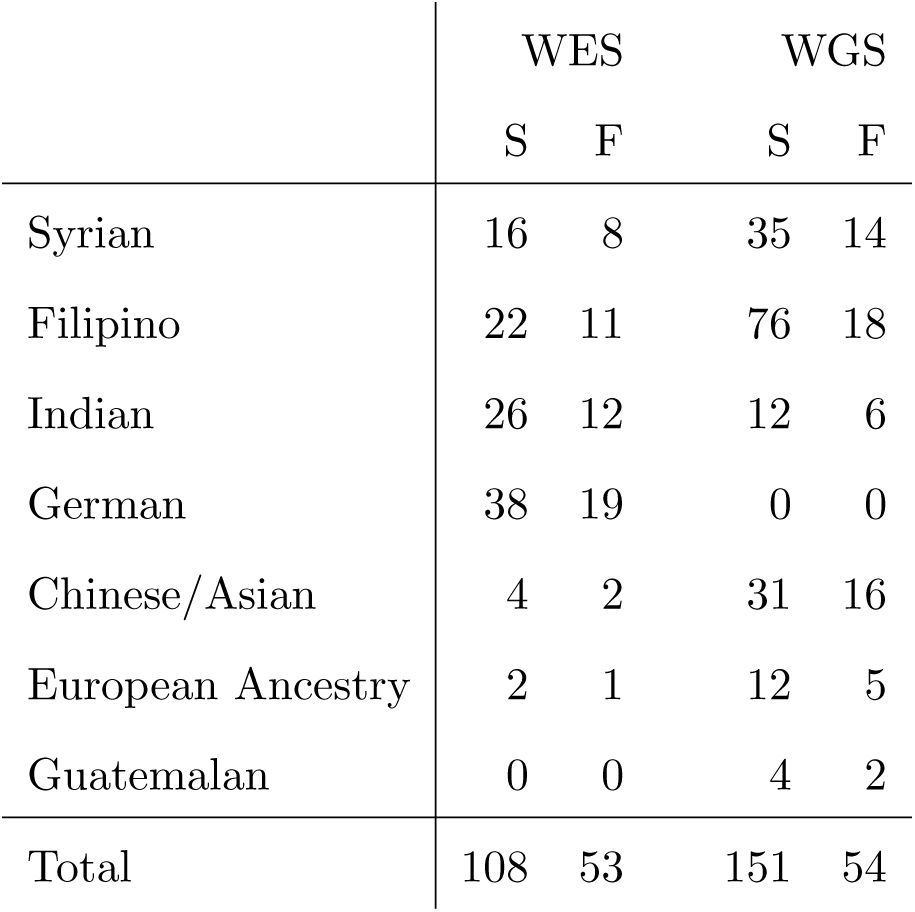
Number of affected individuals and families with nonsyndromic oral clefts, by DNA sequence approach (S: subjects; F: families).

## Results

### Recoding of rare variant haplotypes

It is possible for two or more RVs on distinct haplotypes to be on the same inferred haplotype (due to failure of the inferred haplotypes to distinguish all actual haplotypes), thus being incorrectly recoded together as one RV. To assess the frequency of such an event, we applied our RV haplotype recoding procedure to intervals defined from 5 kb upstream to the transcription end of each RefSeq gene (hg19 assembly) in a dataset simulated as described in the “Simulation study” section above. There were 18,272 genes with at least one coding RV in the CEU, TSI and GBR samples. Taking the original estimated haplotypes as the truth and processing each family separately, we generated a total of 450,114 family-specific haplotypes with at least one coding RV (an average of 24.6 haplotypes per gene, or 0.54 per gene per family). For 99.7% of these RV-bearing haplotypes, all coding RVs recoded to that haplotype were on the same actual haplotype.

### Type I error, power, and scalability

We computed the potential p-values as defined by Bureau et al. [2014b] for every replicate at every penetrance level. These potential p-values were generally less than 10^−5^, and thus the simulated datasets were sufficiently informative to potentially reject the null hypothesis at the adopted significance levels. In the simulation under the null hypothesis, computation of the partial sharing gene-based test succeeded in 97% of 1000 replicates (those where RVs were seen in 10 or fewer families), and computation of the complete sharing test always succeeded in this simulation. Type I error was generally well controlled for RareIBD, the two versions of the RVS approach, and GESE when the MAF was correctly estimated. A slight inflation at the most extreme p-values was observed however, particularly for GESE (Figure 1). Type I error inflation was severe for the pVAAST CLRTp test, and somewhat substantial for GESE and pVAAST LOD when the MAF was underestimated by a factor of 10.

The performance of the gene-based tests at detecting causal RVs are compared for *PEAR1* (Figure 2, left) and *CDH13* (Figure 2, right). Computation of the subset sharing gene-based test succeeded in 96% and 70% of replicates (where RVs were seen in 10 or fewer families) for *PEAR1* and *CDH13*, respectively. Since for pVAAST 10,000 simulations under the null did not allow us to estimate p-values below *α* = 1 × 10^−5^, we compared power of all methods at significance level *α* = 1 × 10^−4^. In addition, Supplementary Figures 1 and 2 show the power at significance level *α* = 1 × 10^−5^ for methods conditioning on the presence of at least one variant in a gene, and for GESE the more stringent *α* = 2.5 × 10^−6^ level, as suggested by Qiao et al. [2017] to correct for the total number of genes in the genome. Similarly, we divided the significance level by 4 for the stringent GESE (Figure 2 with *α* = 2.5 × 10^−5^). Generally, all tests performed similarly, except the LOD test of pVAAST. RareIBD had in most instances slightly higher power than its nearest competitors. The test allowing for sharing by a subset of subjects had slightly greater power than the complete sharing test for the small *PEAR1* gene and slightly lower power than the complete sharing test for the larger *CDH13* gene. GESE had lower power than RareIBD and the RVS tests when applying the stringent significance level suggested for GESE by Qiao et al. [2017], for a fair comparison with tests conditioning on the presence of at least one variant. Only when using the same *α =*1 × 10^−5^ significance level as the other tests did GESE have slightly higher power in the small *PEAR1* gene (Supplementary Figure 1). The LOD test of pVAAST had low power, with the LOD score maximizing at 0 for all replicates in the small *PEAR1* gene and a majority of replicates in the larger CDH13 gene. When the relative risk of causal RVs was equal to 10 and thus many unaffected carriers are expected, the linkage LOD test of pVAAST had higher power than the other methods in the large CDH13 gene, but this power did not increase further at higher relative risks. The CLRTp combining linkage and case-control association signals gave the highest power under all alternative models for both genes, which is due to additional information beyond variant segregation within families (examined here). Computing times for testing coding RVs in the gene CDH13 in one replicate of the simulated dataset exhibited dramatic differences in scalability (Table 3), with the RVS complete sharing test taking the least computing time (even including the prior haplotype recoding) followed by GESE and the RVS partial sharing test. We also examined the correlation of the − log_10_ p-values of the five tests and found the partial and complete sharing tests, GESE and RareIBD to be highly correlated, while these four tests are only weakly correlated to the pVAAST LOD test (Supplementary Figure 3, illustrated using gene CDH13 with relative risk 20).

**Figure 2:**
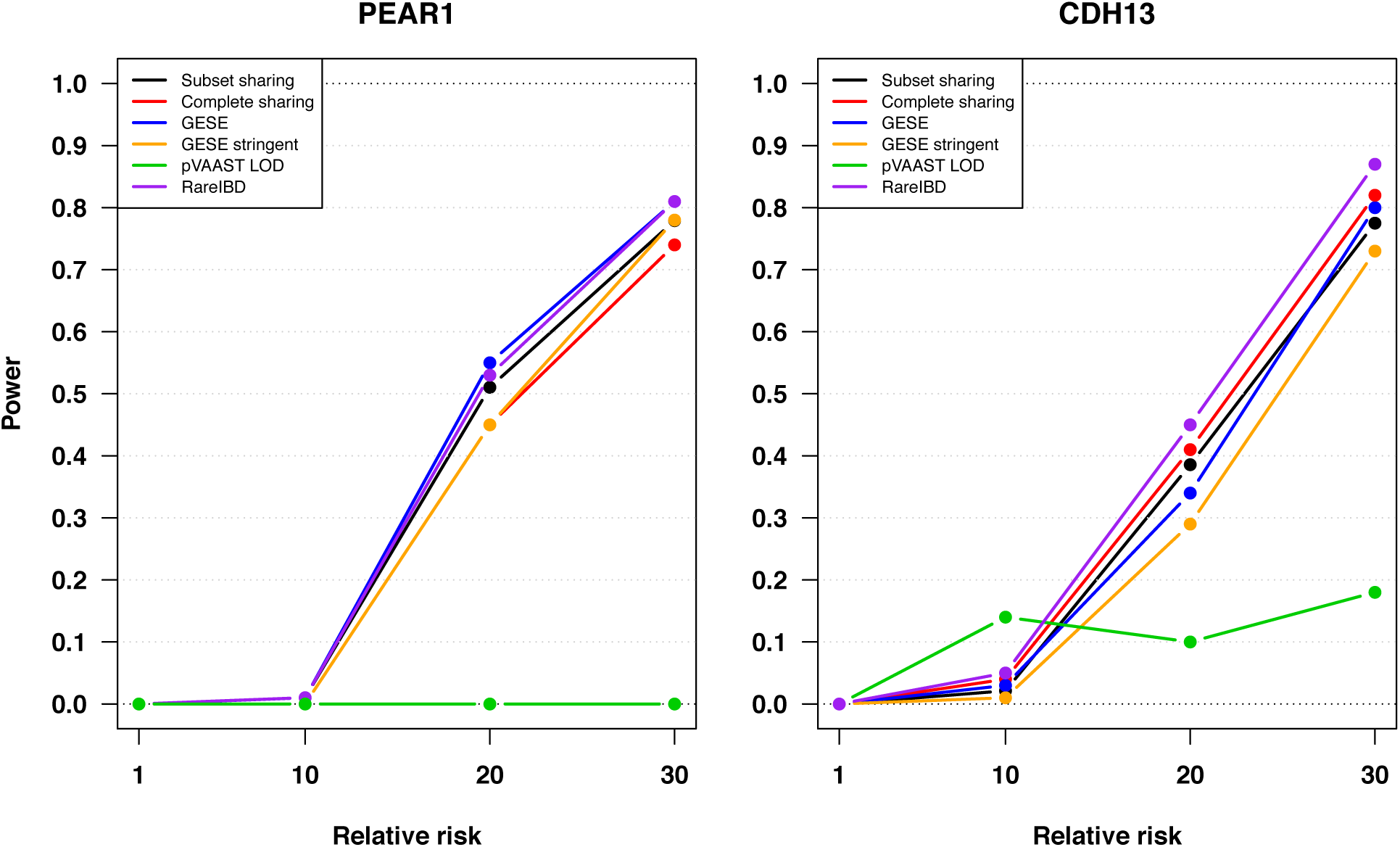
Power for genes PEAR1 (left) and CDH13 (right) under a genetic heterogeneity model with significance level *α* = 10^−4^.

**Table 3:** Running times (in seconds) for analyzing rare coding variants in the gene *CDH13* in one simulated dataset, using a single 2.8 GHz Intel E7-4870 processor of a HP DL580R07 computer. For RareIBD and GESE, the number of resampling simulations was 10^7^. For pVAAST, the number of Monte Carlo simulations to estimate the p-value was 10,000.

| Test | Time [ sec ] |
| --- | --- |
| RVS: haplotype recoding | 2 |
| RVS: complete sharing | 1 |
| RVS: partial sharing | 37 |
| GESE | 18 |
| RareIBD | 2,392 |
| pVAAST | 20,617 |

### Analysis of the oral cleft exome sequence data in the WGS sample

There were 69,719 rare exonic and splice site SNVs passing filtering criteria. The number of genes with rare SNVs for which the RVS p-value computation succeeded in the WGS dataset was 12,706 for the partial, and 14,050 for the complete sharing test. We calculated Bonferroni-corrected significance thresholds using the number of genes with RVs for which the potential p-value as defined by Bureau et al. [2014b] would remain below 0.05 after applying the correction, and obtained 6,372 genes with potential p-values below the threshold 0.05/6,372 = 7.8 × 10^−6^ for a family-wise Type I error rate of 0.05 under the partial sharing test, and 7,647 values below the threshold 0.05/7,647 = 6.5 × 10^−6^ for a family-wise Type I error rate of 0.05 under the complete sharing test.

**Table 4:** The three most significant genes based on the partial sharing test applied to exonic SNVs with frequency ≤ 1% found in the whole genome sequence data of 151 relatives from a sample of 54 multiplex oral cleft families.

| Gene | Based on pedigree structure |  | Adjusted for unknown relationships |  |
| --- | --- | --- | --- | --- |
|  | Partial sharing | Complete sharing | Partial sharing | Complete sharing |
| EP400NL | $4.8 \times 10^{-6}$ | $1.8 \times 10^{-6}$ | 0.0028 | 0.0038 |
| SPATA21 | $2.3 \times 10^{-5}$ | $3.6 \times 10^{-6}$ | $3.7 \times 10^{-5}$ | $8.0 \times 10^{-6}$ |
| IL17RE | $7.6 \times 10^{-5}$ | 0.037 | 0.0015 | 0.0540 |

Among the top three genes based on the partial sharing test (Table 4), E1A binding protein p400 pseudo-gene 1 (*EP400NL*) and spermatogenesis associated 21 (*SPATA21*) were also the top hits with the complete sharing test, which returned smaller p-values than the partial sharing test. In both genes, shared variants appear in families of Syrian origin, where unknown relationships are likely. We adjusted the RV sharing probabilities based on a mean kinship of 0.013 among founders, estimated from the Syrian families of the WES sample as we did previously [Bureau et al. 2014b]. This increased the sharing probabilities above the Bonferroni corrected significance threshold. The partial sharing test also detected the gene interleukin 17 receptor E (*IL17RE*), with eight families carrying a RV (Supplementary Figure 5). Four of these eight families were Syrian, and the adjustment for unknown relationships performed as before increased the p-value to 0.0015. Sharing by two or more subjects was observed in five families, but sharing by all affected relatives occurred in only two families, so the complete sharing p-value was much higher. *IL17RE* (OMIM: 614995) appears to play a role in host mucosal defense against infection. Some evidence of association between common SNVs in *SPATA21* and cleft lips with or without cleft palate (*p* = 2.9 × 10^−4^) was reported in a multiethnic genome-wide association study [Leslie et al. 2016]. No evidence of association was reported for *EP400P1* and *IL17RE* in any study included in the Facebase database.

### False signals due to sharing IBS without IBD in the WES sample

In the WES sample, we did not phase rare SNVs with common variants since we only had the exome sequence, and relied solely on filtering by variant frequency in the 1000 Genomes, Exome Sequencing Project and gnomAD (exome data) databases. Where there were multiple RVs in the same gene in a family, we retained the minimal RV sharing probability, effectively merging RVs with the lowest sharing probability. Excess RV sharing was detected in the gene *ADAMTS9* using the complete sharing test. A supposedly rare SNV was present in at least one affected individual from 26 families: the variant was shared by the two affected subjects in seven families while the variant was seen in only one out of two affected subjects in 18 families and in only one out of three affected subjects in one family (*p* = 4.0 × 10^−6^). Upon inspection of all SNVs from the genotyping array and Merlin-inferred haplotypes in *ADAMTS9*, there was evidence against SNVs being IBD between the affected relatives in five of the seven families where the variant was shared IBS (due to homozygous genotypes for opposite alleles in two affected relatives), including the three families where Bureau et al. [2014b] reported the G allele at rs149253049 was shared by two affected relatives.

### Rare SNVs in WES sample within genes with a signal in the WGS sample

We examined rare SNVs found in the WES sample within genes reported in Table 4. There were two rare SNVs in *EP400NL*, each seen in a single individual. There were two SNVs in *SPATA21* shared by two affected relatives (in two separate families), one of these two inferred IBD and the other not. Finally, a SNV in *IL17RE* shared IBS by two out of three relatives in one family could be inferred not to be IBD.

## Discussion

This article introduced a number of improvements to the RVS approach for linking RVs to disease introduced by Bureau et al. [2014b]. The approach can now be applied to a gene-based analysis, the commonly used strategy to jointly test all RVs in a gene and increase the number of variant carriers [Li and Leal 2008, Lee et al. 2014]. This requires a strategy to work with multiple RVs occurring in the same family. When genomic sequence is available, inferring haplotypes of common and previously known RVs within any one gene enables identification and merging of all RVs on the same haplotype prior to analysis, and avoids recomputation of the same RV sharing probability for each of these RVs. There may still remain RVs on different haplotypes within the same family. This is a rare occurrence even in genes with multiple RVs such as *CDH13* used in our simulation, and taking the minimal sharing probability among them did not lead to inflated Type I errors in our simulation study.

Phenocopies, diagnosis errors and intra-familial genetic heterogeneity clearly exist, so causal RVs may not be shared by all affected relatives in any one pedigree. One way to detect excess sharing among sequenced family members is then to examine partial sharing, i.e. sharing by a subset of affected subjects in a family, and summing the probability of sharing patterns as or more extreme as the observed one within and between families when computing the p-value. The test allowing partial sharing had a slight power advantage over the complete sharing test for a gene with few RVs in our simulation based on the oral cleft families, but a slight power disadvantage for a gene with many RVs, despite simulating data under genetic models with phenocopies. This difference can be explained by an increased chance of partial sharing under the null hypothesis when there are many RVs, while sharing by all affected subjects remains rare. The exponential growth in computational complexity of the partial sharing test p-value calculation with the number of affected subjects in families with a RV restricts the size and number of families included in the analysis. Nonetheless, the partial sharing test may detect excess RV sharing patterns not detected by the complete sharing test, as exemplified by the sharing pattern observed in the *IL17RE* gene in our WGS sample (although cryptic relatedness in the population increased the p-value to a non significant level, so the observed sharing patterns may well result from RVs not involved in susceptibility to oral cleft).

Another way to detect excess sharing without requiring all affected subjects to share a RV is the RareIBD approach based on the number of affected subjects carrying a RV (and unaffected subjects not carrying the RV when such subjects are available). The RareIBD and RVS tests coincide in the special case where all pedigrees have only one affected relative pair of the same type (e.g. an affected first cousin pair), founders are not sequenced, and there is at most one RV per family in a given gene. The RareIBD score can then only take two values (2 or 1) in all families where a RV is seen, the same standardization of that score is applied in all these families [Equation 4 of Sul et al. 2016], and all families have the same weight in the weighted summation over families [Equation 5 of Sul et al. 2016]. The RareIBD Z-score is thus a linear function of the number of families where the score is 2, which under the null hypothesis follows a binomial distribution with success probability being the probability of sharing a RV by both affected relatives given at least one is a carrier (e.g. 1/15 for affected first cousin pairs). In this setting both RVS tests simplify to the exact tail probability of the same null binomial distribution (and while our method provides a closed-form solution, the gene-dropping simulation used in RareIBD is sampling from the binomial distribution to compute the p-value). When all pedigrees have the same configuration of three or more affected subjects, there will be slight differences between RareIBD and each of the RVS tests due to differences in the ordering of the combinations of complete and partial sharing events across families, and hence of their null distributions. Nonetheless, the most extreme event where all affected subjects in all families share a RV has the same p-value, because the p-value is then again the probability of that most extreme event, whether it is computed exactly or by gene dropping. When different configurations of affected subjects are found in different pedigrees, as in our simulated pedigree sample, standardization and weighting of the pedigrees in Rare IBD introduces further differences with the RVS tests. The correlation between the −*log*_10_*p* of the RareIBD and each RVS test remains high, as evidenced in Supplementary Figure 3, but this figure also reveals datasets simulated with causal RVs where the RV sharing tests rejected the null hypothesis at a given significance threshold while RareIBD failed to reject it, and vice-versa. Computational complexity of evaluating the RareIBD statistic is linear in the number of subjects once the expectation and standard deviation of the number of subjects sharing a RV have been precomputed, but must be repeated on a large number of replicates of gene-dropping under the null hypothesis to obtain a p-value. As a result, for a dataset of typical size like our simulated dataset of 33 extended families, the time to compute the exact p-value of the partial sharing test can be substantially shorter than the time to compute the RareIBD p-value by gene-dropping simulations (Table 3).

The pVAAST approach combines the case-control and pedigree-based information in a single statistic (CLRTp), which achieved greater power than all other methods assessed (although the pVAAST linkage LOD score alone had low power). However, the CLRTp statistic is highly sensitive to inflation due to variant frequency underestimation, as it creates a frequency difference between the cases and controls captured by the CLRT part of this statistic. The LOD score and, in a different way, the GESE statistic also involve the variant frequency and are inflated to a lesser extent by its underestimation. When families in a sample come from various populations, as was the case with the oral cleft families, methods that do not need variant frequencies can be validly applied to the whole sample, provided the analyzed variants are rare or absent in all populations of origin of the families, while methods requiring variant frequency estimates need to be applied separately in each population or use variant frequencies that are incorrect for most families if applied to all families together. Methods requiring variant frequency estimates also have the disadvantage that all genes with RVs seen in the control sample or reference database must be considered in the multiple testing correction, instead of only those genes with RVs seen in affected subjects. This reduces power, as exemplified in Figure 2 when a significance level 4 times lower was used following the suggestion of Qiao et al. [2017].

All variant sharing methods we evaluated show good power to detect genes containing causal RVs with high relative risks (20 or higher) but much lower power when the relative risk is smaller at the sample size used in our simulation study. It is plausible that very rare variants with high relative risks remain to be discovered for complex traits because family-based and case-control association studies cannot detect variants carried by a very small number of affected subjects. Even if most undiscovered variants have relative risks at which power of variant sharing methods is fairly low, if there are many such variants the power to detect at least one of them can be good.

The second purpose of inferring haplotypes in the vicinity of a putative rare variant is to confirm the IBD status of the variants, and hence their rarity in the population. While no estimate of variant frequency is needed to compute RV sharing probabilities, the inference is nonetheless sensitive to the actual allele frequency in the population [Bureau et al. 2014b]. Variants rare in public databases of diverse human populations (such as the 1000 Genomes or gnomAD) may be not so rare in specific populations. For instance, observing the G allele at rs149253049 on six distinct haplotypes in as many affected subjects from three multiplex families suggested this allele is fairly common in the Bengali population from which these families were sampled. Placing RVs on haplotypes and keeping in the analysis only those found on a single haplotype within a pedigree, as we proposed here and applied to the WGS sample, increases our confidence that these RVs are indeed shared IBD. This is most feasible with whole genome sequence, where all variants are detected with the same technology. Phasing RVs from exome sequence data with SNVs from genotyping arrays would pose a challenge, and we did not implement such a procedure.

Haplotypes of known variants do not perfectly distinguish all haplotypes, and may lead to merging RVs actually on distinct haplotypes. Our assessment by simulation revealed a small 0.3% probability of such events occurring. Also, there may be recombination events occurring within a gene in recent generations resulting in IBD copies of a RV on distinct haplotypes spanning a single gene. Future development will include an improvement of our approach to identify recent recombination events, and define haplotypes over the intervals between recombination events instead of only relying on gene boundaries.

Software user-friendliness is an important factor driving the adoption of methods in statistical genetics. Our RV sharing tests and GESE are available as R packages, and interface with data classes for pedigrees and genotype data defined in the R statistical environment (cran.r-project.org). Users familiar with this environment already know how to use functions from other R packages to read data in the most common format such as variant call format (VCF) and ped, and convert them to these data classes. RareIBD and pVAAST are available as stand-alone software packages, and require data input files in their own format. External scripting or calls to provided formating scripts are needed to reformat data files, which may represent important hurdles for some users. The handling of missing genotypes also distinguishes the evaluated RV analysis packages: our methods and GESE focus on RVs seen in the sequence with missing genotypes treated as the reference allele homozygous genotype, while pVAAST sums over all possible genotypes and RareIBD requires complete genotype data, so missing genotypes must be imputed externally prior to calling the program.

The example of application of RVS to a familial study of oral cleft was limited to the exome. RVS can also be applied to whole genome sequence when it is available, as in one of the two familial samples presented here, to test rare variants in the non-coding part of the genome, where a large proportion of GWAS signals are located. Whole genome analysis presents challenges such as defining meaningful analysis windows outside of genes and selecting variants more likely to be functional based on biological annotations which will be addressed in future work.

In conclusion, the additional features of a gene-based analysis of RV sharing with confirmation of IBD status of variants by inferring haplotypes from genomic sequence data does enhance the ability of our RV sharing approach to detect causal RVs in studies of extended multiplex families, while maintaining its original advantage of not requiring variant frequency estimates from a population sample. Testing for partial sharing patterns offers an additional option to detect RVs involved in complex diseases where phenocopy, mis-diagnosis or intra-familial genetic heterogeneity exist, but this approach did not always provide a power gain over computationally cheaper alternatives.

## Acknowledgements

We thank the members of the families who participated in the oral cleft sequencing studies, and the field and laboratory staff who made this analysis possible. We thank Thomas Sherman (Johns Hopkins Bloomberg School of Public Health) for programming the `RVS` package and Saeed Sabbah (Université Laval) for performing the haplotyping of SNVs in the WES sample. The statistical analysis work was partly funded by the Canadian Statistical Sciences Institute through a Collaborative Research Team grant of which AB is the leader. National Institute of Dental and Craniofacial Research grant R03-DE-02579 to IR. The oral cleft sequencing studies were supported by the National Institutes of Health (NIH) (R01-DE-014581, U01-DE-018993, and U01 DE020073 to THB.; R01-DE-016148 and R01-DE-009886 to MLM.; P50-DE-016215 and R37-DE-08559 to JCM). Recruitment of Syrian families was supported by the Intramural Research Program of the National Human Genome Research Institute, National Institutes of Health, USA, and the Ibn Al-Nafees Hospital, Syrian Arab Republic. Additional support from X01HG006177 to THB, MLM, and JCM for whole exome sequencing at the Center for Inherited Disease Research, which is funded through a federal contract from the NIH to Johns Hopkins University (contract no. HHSN268200782096C). The FaceBase database was funded by NIH grant U01-DE024425 to MLM.

