## Supplementary Materials for "Inferring Disease Risk Genes from Sequencing Data in Multiplex Pedigrees Through Sharing of Rare Variants" (from another version of this preprint: version 2, 2018-04-04)

#### Supplemental data

**Alexandre Bureau[1,2*], Ferdouse Begum[3], Margaret A. Taub[4], Jacqueline Hetmanski[3], Margaret M. Parker[5], Hasan Albacha-Hejazi[6], Alan F. Scott[7], Jeffrey C. Murray[8], Mary L. Marazita[9], Joan E. Bailey-Wilson[10], Terri H. Beaty[3], Ingo Ruczinski[4]**

[1] Département de Médecine Sociale et Préventive, Université Laval, Québec, Canada.

[2] Centre de recherche CERVO, Québec, Canada.

[3] Department of Epidemiology, Johns Hopkins Bloomberg School of Public Health, Baltimore, MD, USA.

[4] Department of Biostatistics, Johns Hopkins Bloomberg School of Public Health, Baltimore, MD, USA.

[5] Channing Division of Network Medicine, Harvard Medical School, Boston, MA, USA.

[6] Prime Health Clinic Jeddah, Riyadh, Saudi Arabia.

[7] Institute of Genetic Medicine, Johns Hopkins Medical Institutions, Baltimore, MD, USA.

[8] Department of Pediatrics, School of Medicine, University of Iowa, Iowa City, IA, USA.

[9] Department of Oral Biology, School of Dental Medicine, University of Pittsburgh, Pittsburgh, PA, USA.

[10] Inherited Disease Research Branch, National Human Genome Research Institute, Baltimore, MD, USA.

### SUPPLEMENTARY TABLES

| N affected | Freq. | Most distant relatives | kinship | Freq. |
|---:|---:|---:|---:|---:|
| 2 | 33 | 1st degree | $\frac{1}{4}$ | 14 |
| 3 | 5 | 2nd degree | $\frac{1}{8}$ | 16 |
| 4 | 11 | 3rd degree | $\frac{1}{16}$ | 12 |
| 5 | 4 | 4th degree | $\frac{1}{32}$ | 3 |
| 6 | 1 | 5th degree | $\frac{1}{64}$ | 5 |
| | | $\geq$ 6th degree | $\leq \frac{1}{128}$ | 4 |
| Total | 54 | | | 54 |

**Supplementary Table 1:** Descriptive statistics of the WGS family sample.

### SUPPLEMENTARY FIGURES

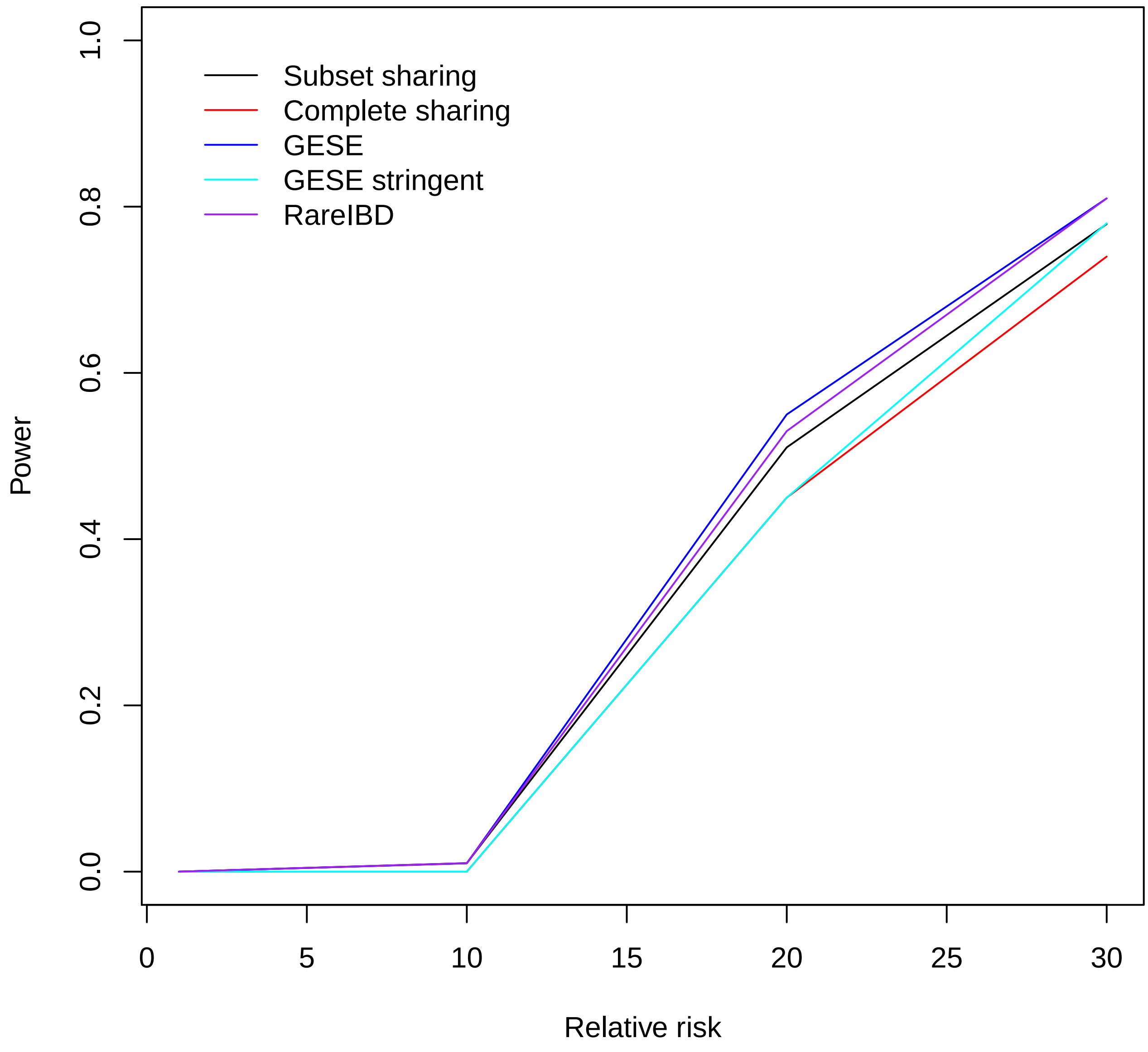


**Supplementary Figure 1:** Power for gene PEAR1 under heterogeneity model at significance level $\alpha = 1x10^{-5}$.

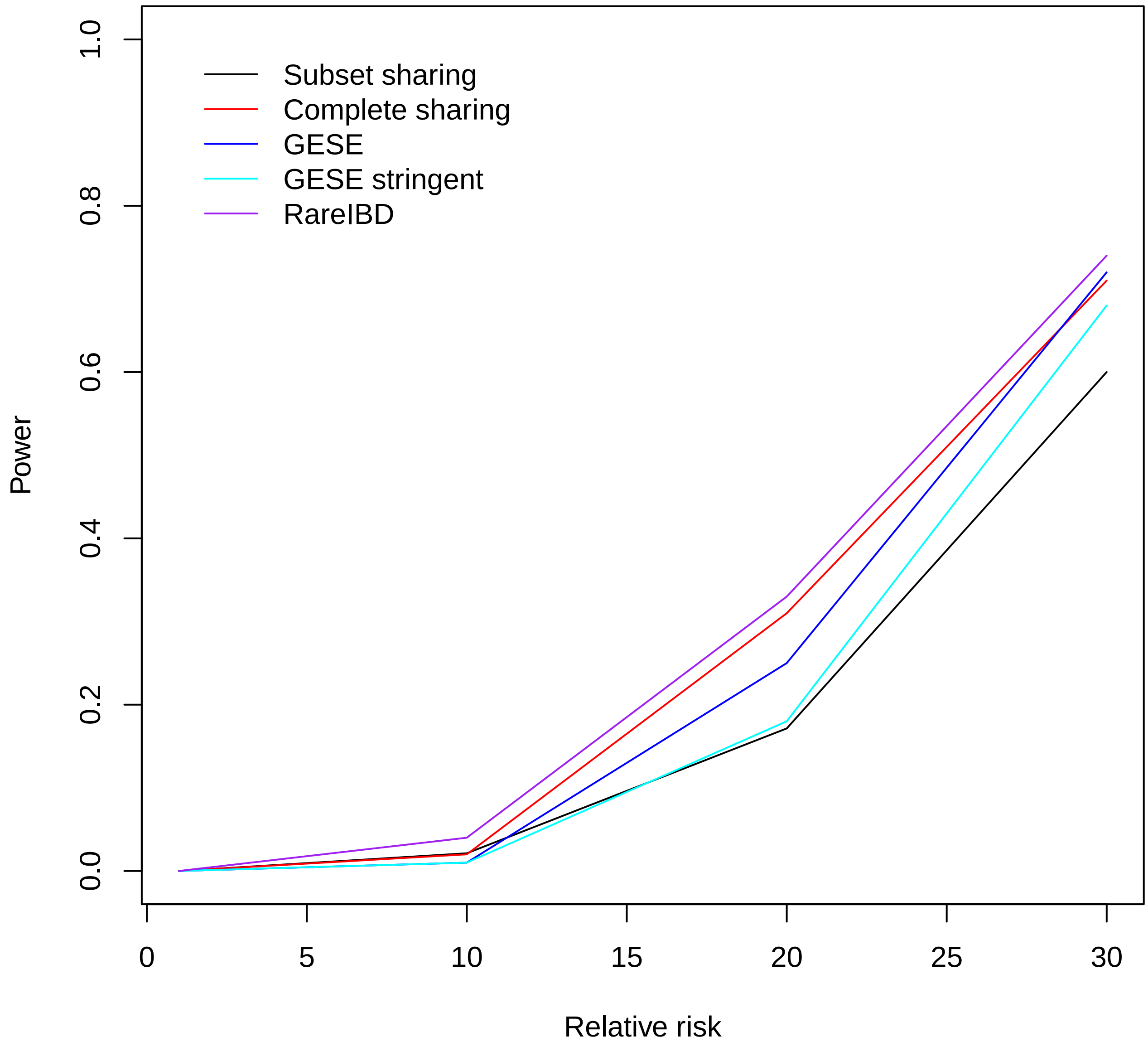


**Supplementary Figure 2:** Power for gene CDH13 under heterogeneity model at significance level $\alpha = 1x10^{-5}$ .

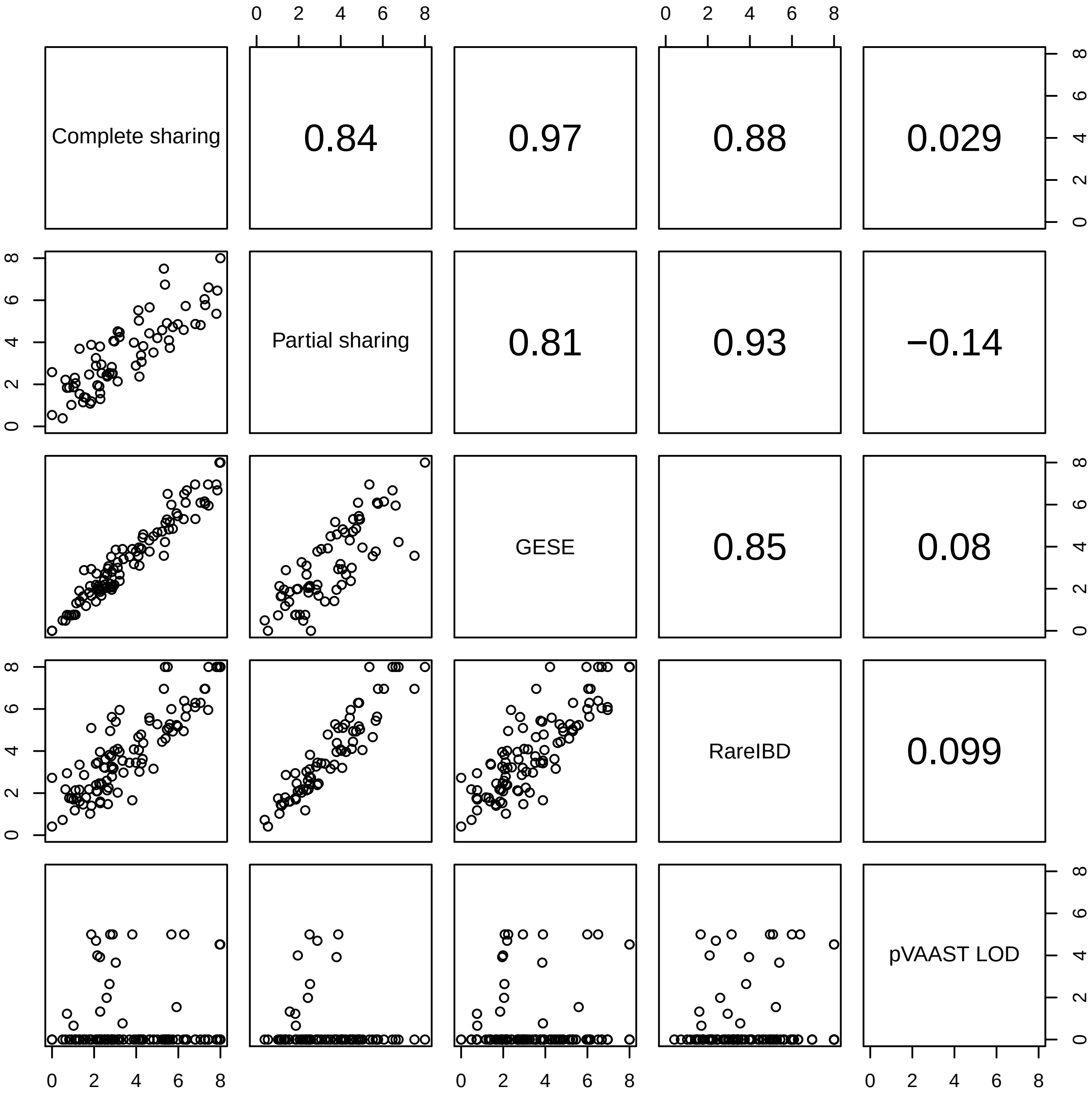


**Supplementary Figure 3:** Comparison of $-\log_{10}$(p-value) of five tests for gene CDH13 when the relative risk is 20. The correlation between the $-\log_{10}$(p-value) is displayed in the upper triangle.

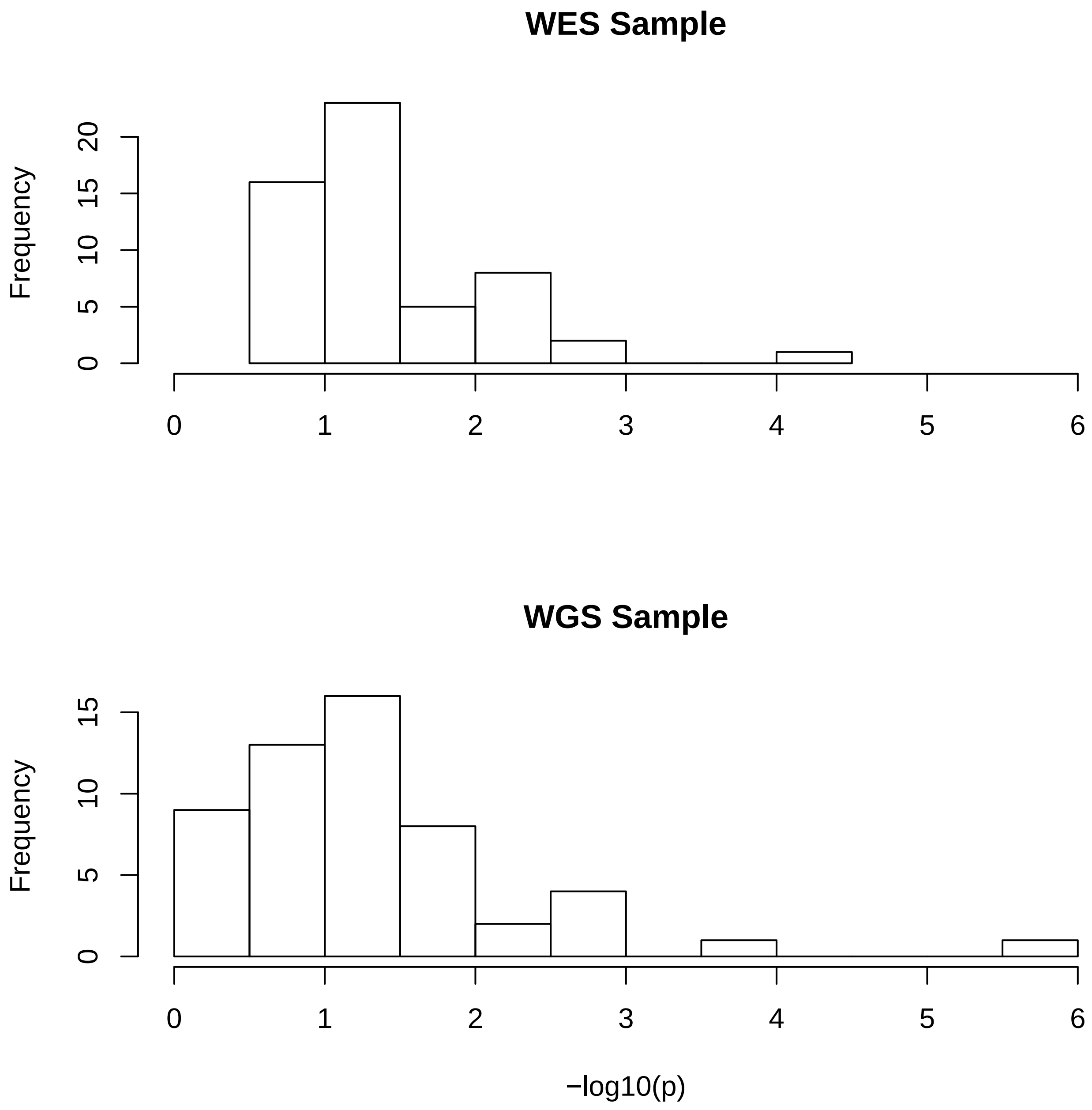


**Supplementary Figure 4:** Distribution of the probability of sharing a rare variants by all affected relatives in a family. Both samples are comprised of multiplex oral cleft families. The WES sample is a sample of 53 families where 2 or 3 distant affected relatives were sequenced. The WGS sample is a distinct sample of 54 families where 2 to 6 close and distant affected relatives were sequenced.

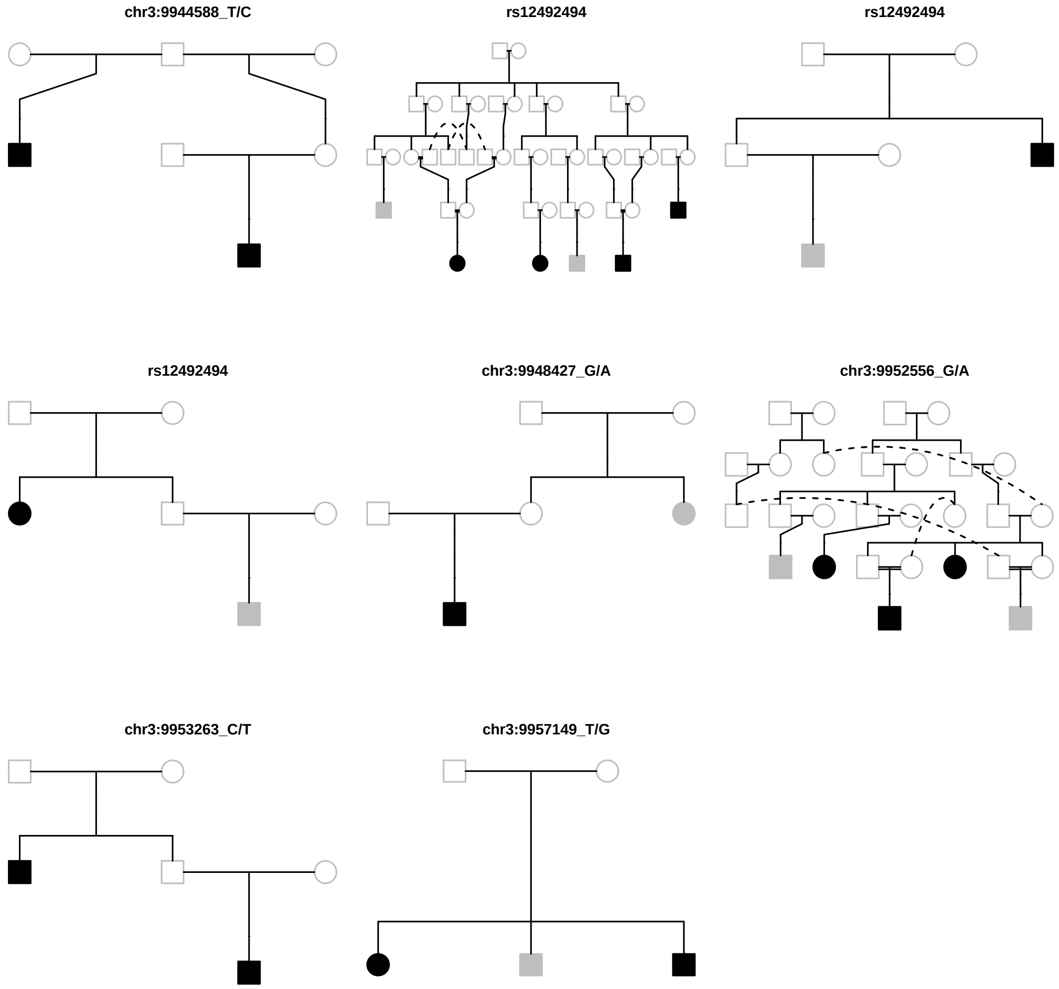


**Supplementary Figure 5:** Families from the WGS sample with an observed RV in *IL17RE*. Black symbols indicate sequenced affected subjects who are carriers of RVs. Grey symbols represent sequenced affected subjects who did not carry any RV. Panel titles indicate rare SNVs reference sequence number or chromosome:position_reference allele/variant allele. Positions are in hg19 coordinates.
